# Na_V_1.7 mRNA and protein expression in putative projection neurons of the human spinal dorsal horn

**DOI:** 10.1101/2023.02.04.527110

**Authors:** Stephanie Shiers, Geoffrey Funk, Anna Cervantes, Peter Horton, Gregory Dussor, Stephanie Hennen, Theodore J. Price

**Affiliations:** University of Texas at Dallas, School of Behavioral and Brain Sciences and Center for Advanced Pain Studies; Southwest Transplant Alliance; Grunenthal Gmbh, Aachen, Germany

**Author notes:** Corresponding author Theodore J Price, University of Texas at Dallas, School of Behavioral and Brain Sciences, 800 W Campbell Rd, BSB 14.102, Richardson TX 75080. GF –. GD –. TJP –.

**Keywords:** Na_V_1.7, spinal cord, dorsal horn

## Abstract

Na_V_1.7, a membrane-bound voltage-gated sodium channel, is preferentially expressed along primary sensory neurons, including their peripheral & central nerve endings, axons, and soma within the dorsal root ganglia and plays an integral role in amplifying membrane depolarization and pain neurotransmission. Loss- and gain-of-function mutations in the gene encoding Na_V_1.7, *SCN9A*, are associated with a complete loss of pain sensation or exacerbated pain in humans, respectively. As an enticing pain target supported by human genetic validation, many compounds have been developed to inhibit Na_V_1.7 but have disappointed in clinical trials. The underlying reasons are still unclear, but recent reports suggest that inhibiting Na_V_1.7 in central terminals of nociceptor afferents is critical for achieving pain relief by pharmacological inhibition of Na_V_1.7. We report for the first time that Na_V_1.7 mRNA is expressed in putative projection neurons (NK1R+) in the human spinal dorsal horn, predominantly in lamina 1 and 2, as well as in deep dorsal horn neurons and motor neurons in the ventral horn. Na_V_1.7 protein was found in the central axons of sensory neurons terminating in lamina 1-2, but also was detected in the axon initial segment of resident spinal dorsal horn neurons and in axons entering the anterior commissure. Given that projection neurons are critical for conveying nociceptive information from the dorsal horn to the brain, these data support that dorsal horn Na_V_1.7 expression may play an unappreciated role in pain phenotypes observed in humans with genetic *SCN9A* mutations, and in achieving analgesic efficacy in clinical trials.

## Introduction

In the mid-2000s, it was found that loss- and-gain-of-function mutations in the gene encoding voltage-gated sodium channel (Na_V_) 1.7 (Baker and Nassar, 2020), *SCN9A*, resulted in congenital pain insensitivity (Cox et al., 2006) or in extreme pain disorders like primary erythromelalgia (Yang et al., 2004; Mann et al., 2019), paroxysmal extreme pain disorder (Fertleman et al., 2006; Hua et al., 2022), and idiopathic small fiber neuropathy (Faber et al., 2012), respectively. The rarity of finding a gene that is critical for sensory pathophysiology and that is backed by human genetics framed Na_V_1.7 as a prime therapeutic target for pain treatment. Several Na_V_1.7 inhibitors have been developed, and a series of clinical trials have been conducted with mixed reports of success (Price et al., 2017; McDonnell et al., 2018; Kingwell, 2019; Alles and Smith, 2021; Biogen, 2021; Eagles et al., 2022). While these outcomes could be due to poor drug pharmacokinetics and/or limited selectivity towards other sodium channels, one hypothesis is that peripherally restricted drugs lack significant efficacy because spinal Na_V_1.7 must be targeted to achieve analgesia. This idea has been supported by mouse studies which suggest that Na_V_1.7 in the central terminals of nociceptors is critical for nociceptive neurotransmission in the dorsal horn (MacDonald et al., 2021). However, it is also notable that mouse genetic studies generating gain-of-function mutations in *Scn9a* have failed to recapitulate human phenotypes (Chen et al., 2021).

Na_V_1.7 is predominantly expressed in the peripheral nervous system in sympathetic neurons and nociceptive sensory neurons in the dorsal root ganglia (DRG) and trigeminal ganglia (Hameed, 2019a). Subcellularly, it is localized to the membrane of sensory neurons including their soma, peripheral axons that innervate the skin, muscle and other organs, and central axons that cross the blood brain barrier and terminate in the dorsal horn of the spinal cord (Black et al., 2012; Shiers et al., 2020; Shiers et al., 2021). It is a key regulator of neuronal excitability as it mediates Na currents during membrane depolarization and action potential firing (McDermott et al., 2019; Middleton et al., 2022) and is dysregulated in pathological pain conditions in both rodents and humans (Black et al., 2008; Li et al., 2018; Sun et al., 2018; Hameed, 2019b; Alvarez et al., 2021; Liu et al., 2021; Nascimento et al., 2022). For this reason, the peripheral nociceptive system has been considered as the primary site of action for therapeutic targeting. However, peripherally restricted Na_V_1.7 inhibitors like Pfizer’s small molecule inhibitor, PF-05089771, and Teva/Xenon’s topical small-molecule inhibitor, TV-45070, failed to show significant analgesic efficacy in clinical trials (Price et al., 2017; McDonnell et al., 2018), suggesting that central nervous system-targeting of Na_V_1.7 may be critical for analgesia.

While absent in the rodent brain(Lein et al., 2007; Allen Institute for Brain Science, 2022a) and human cortex,(Allen Institute for Brain Science, 2022b) Na_V_1.7 protein appears to be entirely localized to presynaptic axons and fibers in both the rodent and human spinal dorsal horn (Black et al., 2012; Shiers et al., 2021). A central analgesic mechanism for Na_V_1.7 was recently proposed wherein Na_V_1.7 -null mice that were pain insensitive, retained nociceptor firing properties but displayed opioid-dependent impaired synaptic transmission and neurotransmitter release at sensory neuron presynaptic terminals in the spinal dorsal horn (MacDonald et al., 2021). Indeed, this could be an important mechanism of action for Na_V_1.7’s key role in nociception but there is some evidence, although inconsistent, for Na_V_1.7 expression post-synaptically within the spinal cord. Na_V_1.7 mRNA has been identified in several subsets of mouse spinal cord neurons using single-cell RNA sequencing (Russ et al., 2021), but only its motor neuron expression could be confirmed with *in situ* hybridization (Alles et al., 2020; Allen Institute for Brain Science, 2022a). However, electron microscopy revealed a substantial proportion of Na_V_1.7 immunoreactivity was localized to dendrites of mouse spinal dorsal horn neurons, but these authors proposed that the protein originated from presynaptic fibers and was transferred to post-synaptic sites through a mechanism that remains to be elucidated (Alles et al., 2020).

We hypothesized that *SCN9A* mRNA and Na_V_1.7 protein might be expressed by human dorsal horn projection neurons. If post-synaptic Na_V_1.7 exists, especially within the pain neurocircuitry in the spinal dorsal horn such as projection neurons, this would provide important evidence to explain the lack of efficacy of peripherally restricted Na_V_1.7 inhibitors and invigorate research towards investigating how Na_V_1.7 expression in intrinsic dorsal horn neurons regulates nociception in addition to its well-described role in nociceptors.

## Methods

### Tissue preparation

All human tissue procurement procedures were approved by the Institutional Review Boards at the University of Texas at Dallas. Human lumbar spinal cords were surgically extracted using a ventral approach (Valtcheva et al., 2016) from organ donors within 4 hours of cross-clamp, frozen immediately in dry ice, and stored in a -80°C freezer. All tissues were recovered in the Dallas area via a collaboration with the Southwest Transplant Alliance. Donor information is provided in **Table 1**. All tissues were collected from neurologic determination of death donors. Human spinal cords were gradually embedded in OCT in a cryomold by adding small volumes of OCT over dry ice to avoid thawing. All tissues were cryostat sectioned at 20 μm onto SuperFrost Plus charged slides.

**Table 1.** Donor Demographic Information

| Donor ID | Age | Sex | Race | Cause of Death | Experiment used |
| --- | --- | --- | --- | --- | --- |
| AICS171 | 19 | M | White | Anoxia/Drug Overdose | RNAscope SCN9A/TACR1 |
| AIAG131 | 61 | F | White | Anoxia/Cardiac Arrest | RNAscope SCN9A/TACR1 |
| AHLC292 | 51 | M | White | CVA/Stroke | RNAscope SCN9A/TACR1 |
| AIBR013 | 18 | F | Black | Anoxia/Seizure | IHC Na<sub>v</sub>1.7/CGRP/Bassoon |
| AHJZ471 | 24 | M | White | Head Trauma/MVA | IHC Na<sub>v</sub>1.7/CGRP/Bassoon |
| AHJD193 | 53 | F | White | CVA/Stroke | IHC Na<sub>v</sub>1.7/CGRP/Bassoon |
| AIEJ414 | 69 | M | Black | CVA/Stroke | IHC Na<sub>v</sub>1.7/CGRP/Bassoon |
| AJD2236 | 56 | M | White | Head Trauma/GSW | IHC Na<sub>v</sub>1.7/Ankyrin-G/MAP2 |
| AIFN459 | 44 | F | Hispanic | CVA/Stroke | IHC Na<sub>v</sub>1.7/Ankyrin-G/MAP2 |
| AJEK100 | 22 | M | Black | Overdose/Cardiovascular | IHC Na<sub>v</sub>1.7/Ankyrin-G/MAP2 |
| AIGD395 | 45 | F | Black | CVA/Stroke | IHC Na<sub>v</sub>1.7/Ankyrin-G/MAP2 |

Sections were only briefly thawed in order to adhere to the slide but were immediately returned to the -20°C cryostat chamber until completion of sectioning. The slides were then immediately utilized for histology. 3 sections from each donor were stained in each experiment, and 3-4 donors were used in each experiment. Donor tissues used in each experiment are also indicated in **Table 1**.

### RNAscope *in situ* hybridization

RNAscope (multiplex fluorescent v2 kit, Cat 323100) was performed as instructed by Advanced Cell Diagnostics (ACD). The protease IV digestion (2 minute incubation) was optimized for human spinal cord. The probe combination used was human specific *SCN9A* (Cat 562251, ACD) paired with Cy3 (Cat NEL744001KT, Akoya Biosciences) and *TACR1* (Cat 310701-3, ACD) paired with Cy5 (Cat NEL745001KT, Akoya Biosciences). The third channel (green, 488) was left unstained to detect background lipofuscin. All tissues were checked for RNA quality by using a positive control probe cocktail (Cat 320861, ACD) which contains probes for high, medium and low-expressing mRNAs that are present in all cells (ubiquitin C > Peptidyl-prolyl cis-trans isomerase B > DNA-directed RNA polymerase II subunit RPB1). A negative control probe against the bacterial DapB gene (Cat 310043, ACD) was used to reference non-specific/background label. Tissues were coverslipped with Prolong Gold Antifade Reagent (Cat P36930, Fisher Scientific).

### Immunohistochemistry (IHC)

Slides were removed from the cryostat and immediately transferred to cold 10% formalin (4°C; pH 7.4) for 15 minutes. The tissues were then dehydrated in 50% ethanol (5 min), 70% ethanol (5 min), 100% ethanol (5 min), 100% ethanol (5 min) at room temperature. The slides were air dried briefly and then boundaries were drawn around each section using a hydrophobic pen (ImmEdge PAP pen, Vector Labs). When hydrophobic boundaries had dried, the slides were submerged in blocking buffer (10% Normal Goat Serum, 0.3% Triton-X 100 in 0.1M Phosphate Buffer (PB) for 1 hour at room temperature. Slides were then rinsed in 0.1M PB, placed in a light-protected humidity-controlled tray and incubated in primary antibodies diluted in blocking buffer overnight at 4°C. The next day, slides were washed in 0.1M PB and then incubated in their respective secondary antibody diluted at 1:2000 with DAPI (1:5000; Cayman Chemical; Cat 14285) in blocking buffer for 1 hour at room temperature. The antibodies used are provided in **Table 2**. The sections were washed in 0.1M PB and then covered with True Black (20% diluted in 70% Ethanol), a blocker of lipofuscin, for 1 minute. Sections were then washed in water, air dried and coverslipped with Prolong Gold Antifade reagent (Cat P36930, Fisher Scientific).

**Table 2.** Antibodies used for immunohistochemistry.

| Antibody | Vendor | Catalog # or Clone # | RRID | Dilution/Conc. |
| --- | --- | --- | --- | --- |
| Mouse-anti- Na<sub>v</sub>1.7 | NeuroMab | N68/6 | AB_2184355 | 2µg/mL |
| Rabbit-anti-CGRP | ImmunoStar | 24112 | AB_572217 | 1:1000 |
| Mouse-anti-Bassoon | Enzo Life Sci | SAP7F407 | AB_11181058 | 2µg/mL |
| Mouse-anti-Ankyrin G | NeuroMab | N106/36 | AB_10673030 | 2µg/mL |
| Chicken-anti-MAP2 | Novus Bio | NB300-213 | AB_2138178 | 2µg/mL |
| Goat-anti-rabbit H&L 647 | ThermoFisher | A21245 | AB_2535813 | 1:2000 |
| Goat-anti-chicken H&L 647 | ThermoFisher | A21449 | AB_2535866 | 1:2000 |
| Goat-anti-mouse IgG2a 555 | ThermoFisher | A21137 | AB_2535776 | 1:2000 |
| Goat-anti-mouse IgG2a 488 | ThermoFisher | AB21131 | AB_2535771 | 1:2000 |
| Goat-anti-mouse IgG1 488 | ThermoFisher | A21121 | AB_2535764 | 1:2000 |
| Goat-anti-mouse IgG1 555 | ThermoFisher | A21127 | AB_2535769 | 1:2000 |

### Na_V_1.7 antibody information and validation

The Na_V_1.7 mouse monoclonal antibody has been knockout validated using immunocytochemistry on mouse cultured DRG neurons and using IHC on rat brain (Grubinska et al., 2019). We have also recently shown that this antibody robustly stains human DRG and shows a specific and similar expression pattern to its mRNA (Shiers et al., 2020; Shiers et al., 2021; Tavares-Ferreira et al., 2022).

### Imaging

All sections were imaged on an Olympus FV3000 confocal microscope. Acquisition parameters were set based on guidelines for the FV3000 provided by Olympus. In particular, the gain was kept at the default setting 1, HV ≤ 600, offset = 4, and laser power ≤ 10%. For the MAP2/Na_V_1.7 experiment, an epifluorescent mosaic image of the entire spinal cord section was captured on an Olympus vs120 slide scanner as a means to visualize the precise anatomical location of the confocal images. Laminar boundaries were drawn using a reference atlas (Sengul et al., 2013; Shiers et al., 2021).

For human spinal cord RNAscope, multiple 20X confocal images with overlapping lipofuscin signal were acquired of adjacent regions of the spinal dorsal horn (mosaic imaging). The images were manually stitched together in Adobe Photoshop (v21.2.12, 2020) by overlaying the overlapping lipofuscin in each image. Once the entire spinal dorsal horn was visualized, laminar boundaries were drawn in Adobe Illustrator using a reference atlas (Sengul et al., 2013; Shiers et al., 2021). This imaging was performed on one section from 3 donors.

### Images Analysis

Raw images were brightened and contrasted in Olympus CellSens (v1.18). Images were pseudocolored for visualization purposes.

For RNAscope, the number of nuclei expressing *SCN9A*-only, *TACR1*-only, and *SCN9A*-and-*TACR1* were counted in each laminar region. For density analysis, the number of RNAscope positive nuclei (neurons) was divided by the area of each laminar subregion (neuron / μm^2^). One section from three donors was analyzed. Graphs were generated using GraphPad Prism version 8.4.3 (GraphPad Software, Inc. San Diego, CA USA).

## Results

### *SCN9A* mRNA is expressed in *TACR1*+ putative projection neurons in the human spinal dorsal horn

The substance P receptor (protein: Nk1r, mRNA: *Tacr1*) labels a subset of projection neurons in the rodent spinal dorsal horn that transmit pain and itch information to the parabrachial nucleus in the brainstem (Blomqvist and Mackerlova, 1995; Mantyh et al., 1997; Carstens et al., 2010). These neurons are localized to the superficial laminae, primarily lamina 1 (Mantyh et al., 1997; Carstens et al., 2010), but they are also found in the deep dorsal horn, mostly in lamina V (Brown et al., 1995). We assessed the distribution of *SCN9A* (Na_V_1.7) and *TACR1* (Nk1r) mRNA in the human spinal dorsal horn using RNAscope. We found that *SCN9A* and *TACR1* mRNAs were detected in cells throughout all laminae in the human spinal dorsal horn (**FIG 1A-E**), and in motor neurons in the ventral horn (**FIG 1F**). Large *SCN9A*/*TACR1* co-expressing neurons were prevalent in lamina 1 and were ∼30-40μm in diameter (**FIG 1B**). In the dorsal horn (laminae 1-7), virtually all of the *TACR1*+ cs were *SCN9A*+ (95%), but only 36% of *SCN9A*+ cells were *TACR1*+ (**FIG 1G-H**). *SCN9A*+ and *TACR1*+ co-expressing neurons were more densely populated in laminae 1-2 (**FIG 1I**). This distribution closely resembles recently published human spinal cord spatial transcriptomic data (Yadav et al., 2023) in which both *SCN9A* and *TACR1* mRNAs were found throughout the dorsal and ventral horns (**FIG 2A**). *SCN9A* was detected in a broad population of excitatory and inhibitory dorsal horn neurons using single-nucleus RNA sequencing of human spinal cord (Yadav et al., 2023), many of which co-expressed *TACR1* (**FIG 2B**).

**Figure 1.**
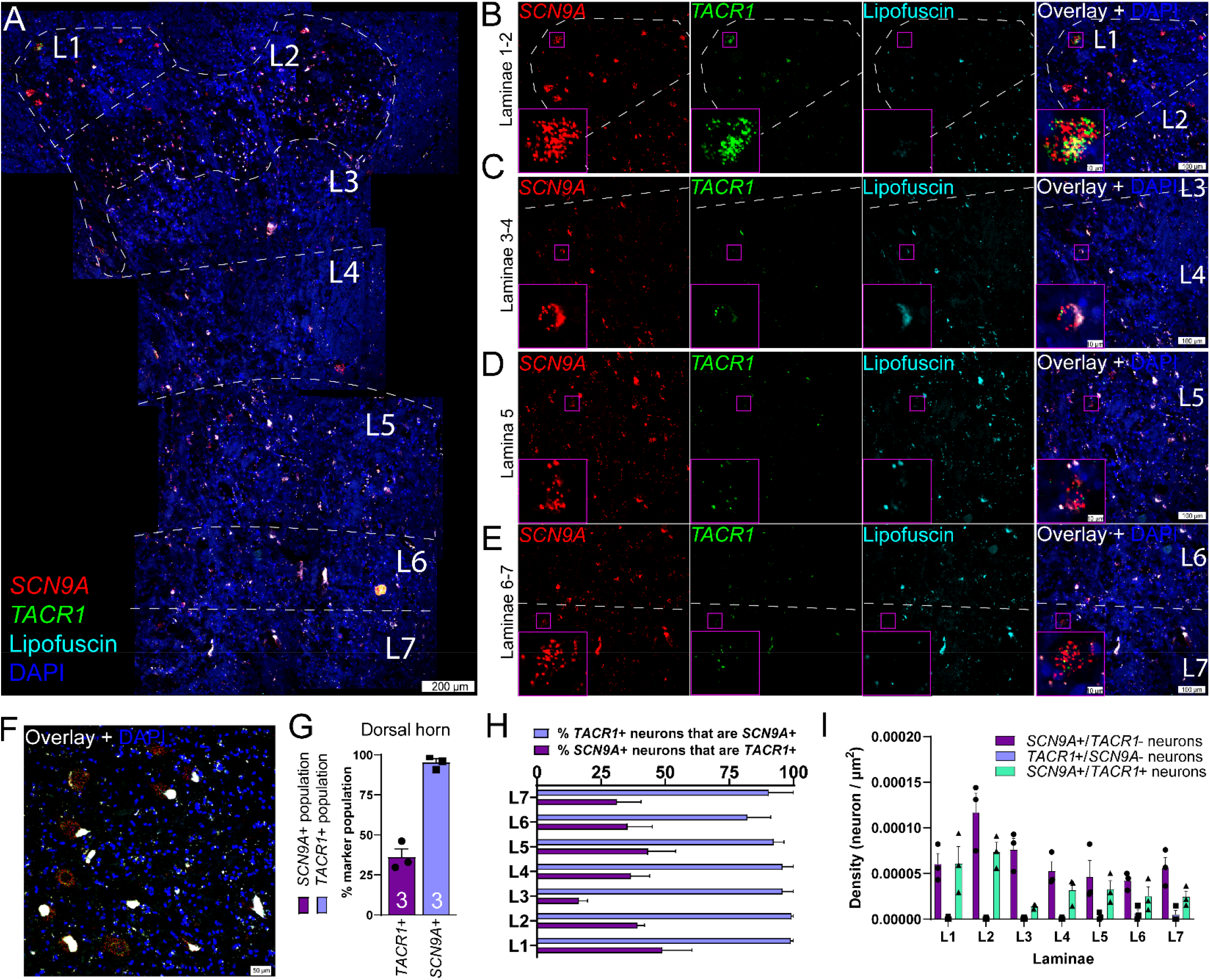
Distribution of *SCN9A* (Na_V_1.7) and *TACR1* (NK1R) mRNAs in the human spinal dorsal horn using RNAscope. **A)** Representative stitched mosaic image of human lumbar spinal cord labeled with RNAscope *in situ* hybridization for *SCN9A* (red) and *TACR1* (green) mRNAs and co-stained with DAPI (cyan). The 488 channel was left unstained (pseudocolored to cyan) to reveal background autofluorescence and lipofuscin which is present in all human neurons. 20X images for each channel are shown for **B)** laminae 1-2, **C)** laminae 3-4, **D)** lamina 5, and **E)** laminae 6-7. The inset images are a zoomed-in, cropped image of a single *SCN9A*/*TACR1* co-positive cell. **F)** *SCN9A* and *TACR1* mRNAs were coexpressed by motor neurons in the ventral horn. **G)** Percentage of *SCN9A*+ neurons in the dorsal horn that coexpressed *TACR1* (purple bar), and the percentage of *TACR1*+ neurons that coexpressed *SCN9A* (blue bar). **H)** Percentage of *TACR1*+ neurons that were copositive for *SCN9A*, and the percentage of *SCN9A*+ neurons that were copositive for *TACR1* for each lamina (L1-L7) of the spinal dorsal horn. **I)** Density of each neuronal subpopulation within each lamina (neuron/μm^2^). Scales bar = panel A: 200 μm; panels B-E: 100 μm and inset is 10 μm; panel F: 50 μm.

**Figure 2.**
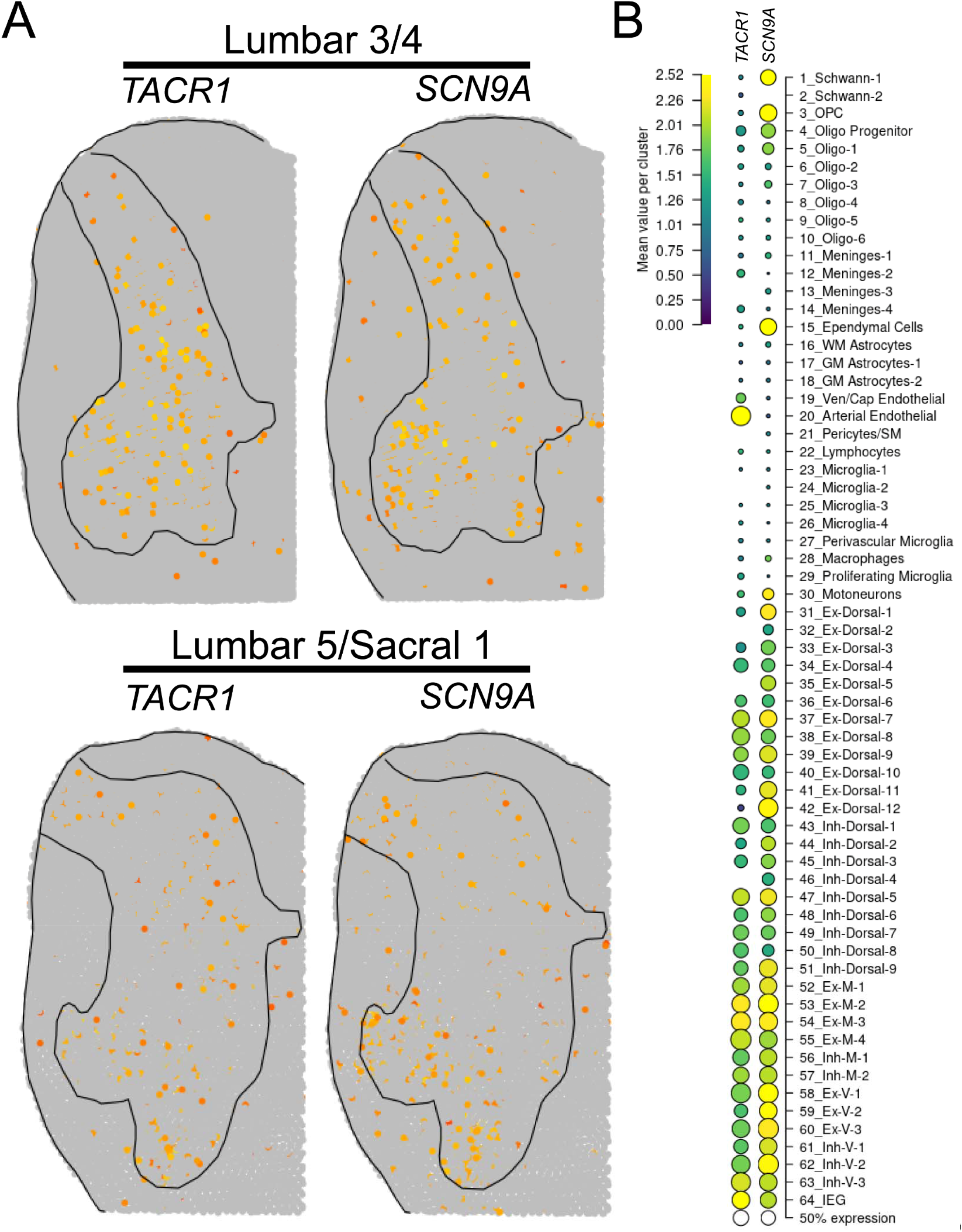
Spatial and single-nuclear RNA-sequencing detection of *SCN9A* and *TACR1* in the human spinal dorsal horn. **A)** Normalized spatial transcriptomic gene expression for *SCN9A* and *TACR1* per barcoded spot on aggregated human lumbar spinal cord sections (Yadav et al., 2023). Solid lines mark grey matter boundaries. **B)** Dot plot showing the average gene expression for *SCN9A* and *TACR1* for each human spinal cord cluster identified using single-nucleus RNA sequencing. Data found at https://vmenon.shinyapps.io/humanspinalcord/

### Na_V_1.7 protein is localized pre-synaptically in the human spinal cord

Na_V_1.7 protein staining in the human spinal cord was robustly found in the dorsal rootlet and substantia gelatinosa which comprises lamina 1 and 2, as we have previously reported (Shiers et al., 2021) (**FIG 3A-3B**).

**Figure 3.**
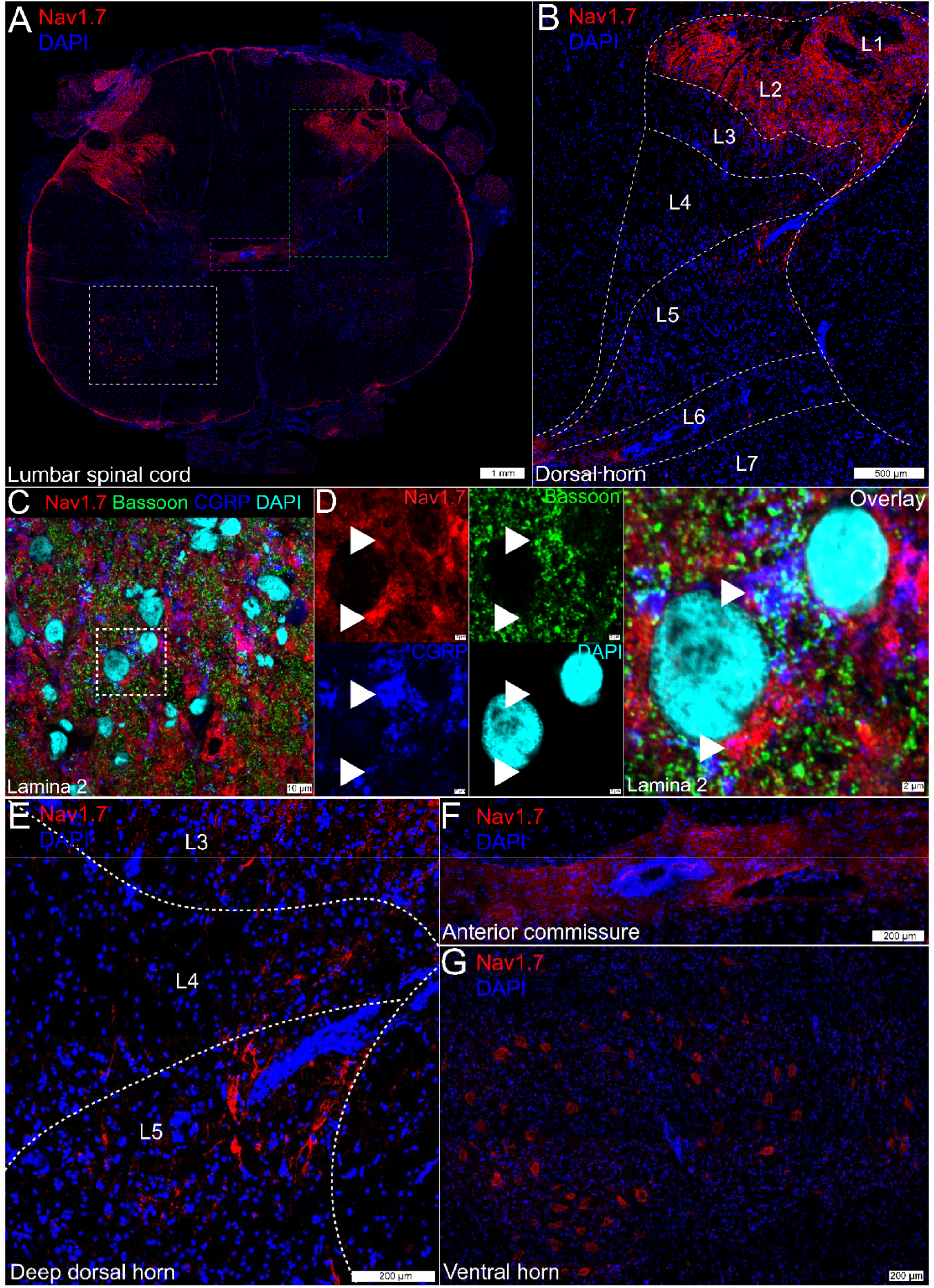
Na_V_1.7 protein expression in the human lumbar spinal cord. **A)** Mosaic image of Na_V_1.7 (red) protein staining in the human lumbar spinal cord, costained with DAPI (blue). The green box represents **B)** the dorsal horn from L1-L6 where robust Na_V_1.7 neuropil staining was observed in lamina 1-lamina 2. **C)** Representative 100X image of Na_V_1.7 (red) protein co-stained with the nociceptive presynaptic marker, CGRP (blue), the presynaptic active zone protein, Bassoon (green), and DAPI (cyan) in lamina 2 of the spinal dorsal horn. A cropped, zoomed in area outlined in white is shown in **D)** where Na_V_1.7 signal is colocalized with CGRP and/or Bassoon (white arrows) or is in close proximity to these proteins. However, not all Na_V_1.7 signal appeared to be localized to the presynaptic compartment. **E)** Axonal Na_V_1.7 was observed in the deeper laminae, most prominently in lamina 5. The magenta box in panel A represents **F)** the anterior commissure where Na_V_1.7 was also detected in axons. The white box in panel A represents **G)** the ventral horn where Na_V_1.7 was localized to the cytoplasm of large motor neurons. Scales bar = panel A: 1 mm; panel B: 500 μm; panels C: 10 μm; panel D: 2 μm; panels E-G: 200 μm.

Na_V_1.7 staining gave a neuropil-like pattern, potentially indicative of synaptic staining, throughout laminae 1-2 (**FIG 3B**). When colabeled with the nociceptive presynaptic marker, CGRP, and the presynaptic active-zone protein, Bassoon (**FIG 3C**), we observed Na_V_1.7 signal that co-localized or was in close proximity to these presynaptic markers (**Fig 3D**). However, not all of the Na_V_1.7 signal in the human spinal cord appeared to be localized to the presynaptic compartment.

Na_V_1.7 was sparsely detected in axons in deeper lamina, with the highest abundance in lamina 5 (**FIG 3E**) and also in axons entering the anterior commissure, where dorsal horn projection neurons cross hemispheres to enter the contralateral spinal thalamic tract (**FIG 3F**). As tracing studies in cats and macaques have demonstrated that neurons projecting to the spinal thalamic tract originate in lamina 1 and lamina 5, it is possible that the non-presynaptic signal in the superficial lamina as well as the axonal signal in the deep lamina and anterior commissure could represent Na_V_1.7 protein expressed by projection neurons (Willis et al., 1979; Jones et al., 1987; Craig, 2006), a hypothesis we explored with additional experiments. Additionally, we observed Na_V_1.7 protein staining in the cytoplasm of motor neurons in the ventral horn (**FIG 3G**).

### Evidence for post-synaptic Na_V_1.7 protein in the human spinal cord

In an effort to identify post-synaptic Na_V_1.7 immunoreactivity in the spinal dorsal horn, we co-stained Na_V_1.7 with a variety of subcellular protein markers. First, we assessed Na_V_1.7 with the cytoskeletal marker, MAP2, which is primarily localized to dendrites (**FIG 4A**). We assessed the dorsal horn from 3 organ donors but observed no convincing evidence for any Na_V_1.7 signal that was localized to the dendritic compartment of resident dorsal horn neurons, nor their soma in neither lamina 1 (**FIG 4B**), nor lamina 2 (**FIG 4C**), nor lamina 4-5 (**FIG 4D**). In the anterior commissure, intensely labeled Na_V_1.7 axonal fibers could be observed crossing hemispheres (**FIG 4E**), but we could not determine what neurons these axons originated from; however, it is likely they are projection neurons as this is a major hub for cross-hemispheric spinal neurotransmission.

**Figure 4.**
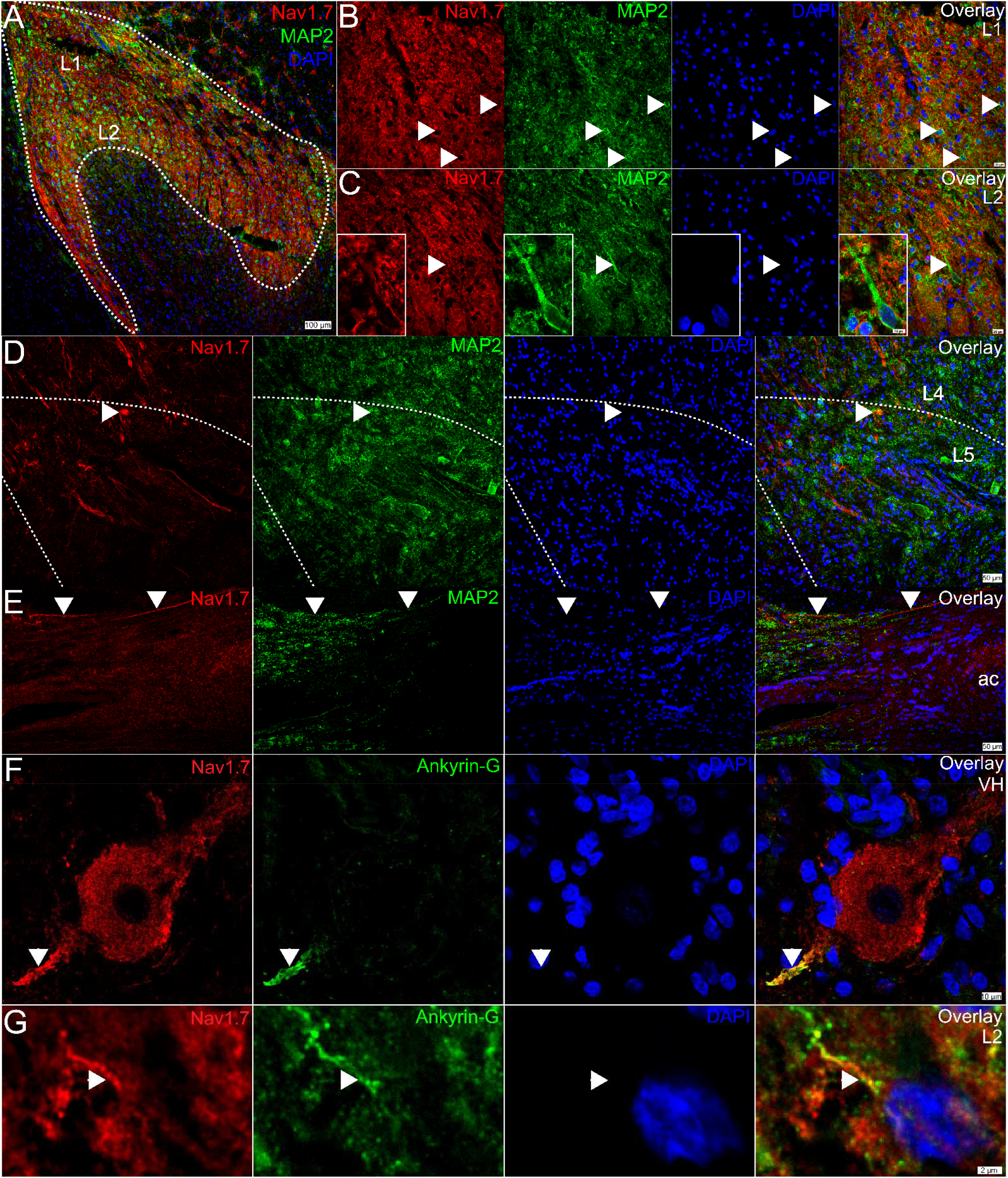
Evidence for post-synaptic Na_V_1.7 expression in the human spinal cord. **A)** Representative 10X image of Na_V_1.7 (red), MAP2 (green), and DAPI (blue) staining in the human lumbar dorsal horn. The white outline demarcates the substantia gelatinosa which comprises lamina 1 and lamina 2. **B)** Representative 40X images of Na_V_1.7 (red), MAP2 (green), and DAPI (blue) in lamina 1, and **C)** lamina 2. White arrows point to MAP2 signal that is localized around resident neurons (appears to be the plasma membrane) but are absent of Na_V_1.7 signal. The image inset in panel C shows a 100X image of a large, L2 neuron with a large apical dendrite that is devoid of Na_V_1.7 signal. **D)** 20X image of Na_V_1.7-positive axonal fibers in the deeper lamina around L4-L5. The white arrow points to Na_V_1.7 and MAP2 copositive signal (yellow in overlay) that does not have a nucleus and is not a cell body. **E)** 20X image of Na_V_1.7 staining in the anterior commissure (ac) where intensely labeled Na_V_1.7-positive axons are highlighted (white arrow). **F)** A 100X image of Na_V_1.7 (red), Ankyrin-G (green), and DAPI (blue) staining in a motor neuron in the ventral horn. **G)** A cropped, zoomed-in image of Na_V_1.7 (red), Ankyrin-G (green), and DAPI (blue) signal in a lamina 2 dorsal horn neuron. Scales bar = panel A: 100 μm; panels B-C: 20 μm; panel C inset: 10 μm; panel D-E: 50 μm; panel F: 10 μm, panel G: 2 μm.

Interestingly, we noted that Na_V_1.7 appeared to be localized to the cytoplasm of motor neurons, including their soma and what appeared to be their axon initial segment (AIS). Recent work with Na_V_1.7 in the rodent DRG found that sensory neurons contain an AIS that it is enriched with Na_V_1.7 and that its localization there is critical for spontaneous activity in neuropathic pain (Nascimento et al., 2022). Other Navs are also known to be enriched in the AIS in the CNS (Leterrier, 2018). To confirm our observation in motor neurons, we co-stained Na_V_1.7 with the AIS marker, Ankyrin-G, and found that, indeed, Na_V_1.7 colocalized with Ankyrin-G at the AIS of motor neurons in the human ventral horn (**Fig 4D**). In the dorsal horn, we also found evidence for Na_V_1.7 localized to the AIS of resident neurons (**Fig 4E**), but this was sparse, and we only observed this pattern in a few neurons across 3 donors. As we see robust Na_V_1.7 mRNA expression in virtually all of the putative (*TACR1*+) projection neurons, its low prevalence in the AIS and absence in the soma and cytoplasm of resident neurons suggests that either Na_V_1.7 is not translated into protein in these neurons, or which is more likely, that it is localized to the membrane of post-synaptic axons. This is further supported by Na_V_1.7 labeling in the anterior commissure.

## Discussion

In the present study, we found evidence for the presence of Na_V_1.7 expression in intrinsic neurons of the human spinal cord, including in the dorsal and ventral horns. First, we identified that *SCN9A* (protein: Na_V_1.7) mRNA was detected in virtually all *TACR1*+ (protein: Nk1r) neurons and that these neurons were more densely populated in lamina 1, 2, 4 and 5, with their lowest density in lamina 3, 6, and 7. In rodents, the Nk1r is expressed by a subset of large-diameter projection neurons that send projections to higher brain regions like the parabrachial nucleus, and are found predominantly in lamina 1 and in the deeper dorsal horn around lamina 5 (Brown et al., 1995; Marshall et al., 1996; Todd et al., 2000). It is important to note that not all rodent projection neurons are *Tacr1*+ (Nk1r+), and interneurons as well as motor neurons express this gene (Russ et al., 2021). Single-nucleus RNA sequencing recapitulates this expression pattern as *SCN9A* and *TACR1* mRNAs were co-expressed in a wide range of human spinal neuronal populations, including motor neurons (Yadav et al., 2023). The high density of the signal in known projection-neuron enriched lamina, as well as the large size profile of the neurons supports that a subset of these *SCN9A*/*TACR1* co-expressing neurons are likely projection neurons.

Similarly, *Scn9a* mRNA was also detected in a variety of *Tacr1* co-expressing neuron populations in mouse spinal cord using single cell sequencing *(Russ et al*., *2021)*. However, only its motor neuron expression could be validated with *in situ* hybridization (Alles et al., 2020; Allen Institute for Brain Science, 2022a). While sensitivity issues could underlie these technical differences, detection of Na_V_1.7 mRNA and protein in rodent spinal dorsal horn neurons has also been inconclusive likely due to its unique subcellular localization to the neuronal membrane which could comprise the soma, dendrites, axon initial segment, nodes of Ranvier, and/or synapse. Indeed, most reports suggest that spinal Na_V_1.7 protein expression is entirely presynaptic due to its robust neuropil staining pattern within lamina 1-2 (Black et al., 2012; Shiers et al., 2021). However, immuno-electron microscopy detected Na_V_1.7 protein localized to dendrites of mouse dorsal horn neurons, but it was hypothesized that these proteins originated in the presynaptic compartment and were transferred to post-synaptic sites through an unknown mechanism (Alles et al., 2020).

While we also detected intense presynaptic Na_V_1.7 labeling in lamina 1-2 of the human spinal cord, not all of its expression was colocalized with presynaptic markers like CGRP and Bassoon. Na_V_1.7 protein was also localized to axons in the deeper lamina, particularly lamina 4 and 5, and to axons in the anterior commissure, the white matter tract connecting the two spinal hemispheres and an important relay for dorsal horn projection neurons transmitting nociceptive information to the contralateral spinothalamic tract (Ku and Morrison, 2022). Because axon tracing methods cannot be employed in the human spinal cord, we could not identify if these were primary afferents or the axons of intrinsic dorsal horn neurons; however, ascending projections of primary afferents ascend in the dorsal columns and do not cross the midline, so it is exceedingly unlikely that these commissural axons are contributed by sensory neurons.

Interestingly, we did not detect Na_V_1.7 localized to dendrites, but instead identified its expression in the AISs of some dorsal horn neurons in lamina1-2 and also in motor neurons. Other Nav family members are highly concentrated at the AIS and their localization there is critical for integration of synaptic currents into action potential generation (Grubb and Burrone, 2010; Leterrier, 2018). Importantly, neurons are known to increase or decrease the size of their AIS as a means to augment or depress their neuronal excitability in response to changing presynaptic input (Grubb and Burrone, 2010; Kuba et al., 2010; Grubb et al., 2011). As sensory neuron hyperexcitability and/or spontaneous activity is a major driver of abnormal nociceptive signaling into the dorsal horn in chronic pain conditions, it is possible that Na_V_1.7 regulation at the AIS in resident dorsal horn neurons is critical for nociceptive processing in the dorsal horn.

While loss-of-function mutations in *SCN9A* result in pain insensitivity in both rodents and humans (Gingras et al., 2014; Shields et al., 2018; Grubinska et al., 2019), gain-of-function mutations engineered in mice to match human mutations that cause pain disorders do not recapitulate the human pain phenotype (Chen et al., 2021). One potential explanation for this is that the loss-of-function phenotype is dependent upon sensory neuron expression of Na_V_1.7 which is conserved across species and the loss of function leads to a conserved loss of action potential generation in nociceptors. On the other hand, the human gain-of-function pain phenotype may require both peripheral sensitization and spinal amplification. As there has been little convincing evidence for Nav1.7 expression in intrinsic neurons of the rodent spinal cord, it is probable that spinal amplification of nociceptive signals does not occur in rodents but may occur in humans with gain-of-function mutations given the broad expression of *SCN9A* in dorsal horn neurons, including putative projection neurons. Gain-of-function mutations in *SCN9A* could lead to increased excitability of projection neurons, which would be further amplified by increased nociceptive input from hyperexcitable nociceptors in the periphery. However, this mechanistic hypothesis may not be as straightforward as this because *SCN9A* mRNA was found in many neurons in the dorsal horn, including inhibitory interneurons by single-nuclear sequencing (Yadav et al., 2023). Nevertheless, dysregulated circuit dynamics in the dorsal horn caused by gain-of-function mutations in Nav1.7 potentially explain the difference in pain phenotype between humans and rodents.

In summary, we offer several compelling pieces of evidence to support the existence of Na_V_1.7 mRNA and protein expression by intrinsic neurons of the human spinal dorsal horn: 1) *SCN9A* mRNA is expressed by human resident dorsal horn neurons detected by both RNAscope *in situ* hybridization, spatial sequencing, and single-nucleus sequencing; 2) all *TACR1*+ human resident dorsal horn neurons express *SCN9A*, a subset of which are likely projection neurons; 3) many *SCN9A*+ and *TACR1*+ co-expressing neurons were large diameter and were most abundant in laminar regions (lamina 1 and lamina 5) that are known to be enriched with projection neurons; 4) not all Na_V_1.7 protein signal was limited to the presynaptic compartment as demonstrated with co-labeling with CGRP and Bassoon; 5) Na_V_1.7 was detected in the axon initial segment of some resident dorsal horn neurons; 6) Na_V_1.7+ axons were detected in the anterior commissure. The existence of Na_V_1.7 in dorsal horn neurons could explain the lack of analgesic efficacy of peripherally restricted Na_V_1.7 inhibitors and offer new insight into a centrally mediated Na_V_1.7 regulatory mechanism for nociceptive processing in the dorsal horn.

## ACKNOWLEDGEMENTS

The authors are grateful to the organ donors and their families for the gift of life and research provided by their organ donation.

